# Differential effects of IL4I1 protein on lymphocytes from healthy and multiple sclerosis patients

**DOI:** 10.1101/2023.09.17.558132

**Authors:** Stephanie E. Davis, Jingwen Hu, Sonia E. Nanescu, Mahesh N. Kumar, Maryna Baydyuk, Helena Oft, Faria S. Amjad, Anton Wellstein, Jeffrey K. Huang

**Affiliations:** Department of Biology, Georgetown University, Washington DC, USA; Interdisciplinary Program in Neuroscience, Georgetown University, Washington DC, USA; Georgetown University School of Medicine, Georgetown University, Washington DC, USA; Department of Neurology, MedStar Georgetown University Hospital, Washington DC, USA; Lombardi Cancer Center, Georgetown University Medical Center, Washington DC, USA

## Abstract

Multiple sclerosis (MS) is a chronic inflammatory disease characterized by immune mediated demyelination of the central nervous system, resulting in extensive neurological deficit and remyelination impairment. We have previously found that interleukin-four induced one (IL4I1) protein modulates CNS inflammation and enhances remyelination in mouse models of experimental demyelination. However, it remained unclear if IL4I1 regulates lymphocyte activity in MS. To assess the therapeutic potential of IL4I1 in MS, we investigated the impact of IL4I1 treatment on human lymphocytes from peripheral blood mononuclear cells (PBMCs) obtained from healthy individuals and MS patients. We found that IL4I1 increased the relative densities of Th2 and regulatory T-cells, while reducing Th17 cell density in healthy control samples. Furthermore, IL4I1-treated lymphocytes promoted CNS remyelination when grafted into demyelinated spinal cord lesions in mice. We found that baseline endogenous *IL4I1* expression was reduced in people with MS. However, unlike healthy controls, IL4I1 treatment had no significant effect on *IL17* or *TOB1* expression in lymphocytes derived from MS patients. These results suggest that IL4I1 skews CD4^+^ T-cells to a regulatory state in healthy human lymphocytes, which may be essential for promoting remyelination. However, IL4I1 appears unable to exert its influence on lymphocytes in MS, indicating that impaired IL4I1-mediated activity may underlie MS pathology.

## INTRODUCTION

Multiple sclerosis is a chronic inflammatory disease, which is characterized by demyelination and neurodegeneration in the CNS.^1–3^ While spontaneous repair of demyelinated lesions is successful during early stages of MS, remyelination ultimately fails in the progressive phase of the disease, leading to axonal dystrophy and irreversible neurological disability.^4,5^ The underlying mechanisms responsible for the transition from successful to failed CNS remyelination are not well understood. It has been proposed that uncontrolled inflammation plays a key role in MS pathogenesis and progression,^6,7^ with autoreactive T cells driving the disease process.^8^ Given the pro-inflammatory skew implicated in MS pathology, immunomodulation becomes a crucial target for promoting the natural repair processes that are deficient in progressive MS.

Interleukin-4 induced protein 1 (IL4I1), a macrophage secreted enzyme,^9–11^ has been shown to modulate inflammation and enhance CNS remyelination in mice by reducing CD4^+^ T-cell driven inflammation.^11^ Additionally, administration of IL4I1 to mice with experimental autoimmune encephalomyelitis (EAE) has been found to promote motor recovery.^11^ In humans, the *IL4I1* gene has been mapped to chromosomal locus 19q13.3-13.4, a region associated with autoimmune susceptibility to lupus, rheumatoid arthritis, diabetes type 1, and MS.^12,13^ Furthermore, it has been suggested that IL4I1 maintains high expression levels of TOB1, an antiproliferative protein that primarily affects Th17 cells.^14^ Notably, expression levels of TOB1 have been inversely correlated with the risk of developing MS from clinically isolated syndrome (CIS).^15^ However, it remains unknown whether IL4I1 affects immune cell activity in MS, or whether altered plasma IL4I1 levels correlate with disease severity or progression. To evaluate its potential as a therapeutic target in MS, here, we investigated the effects of IL4I1 treatment on human lymphocytes derived from PBMCs obtained from healthy individuals and MS patients.

## METHODS

### Patient groups

All human subject work was conducted in accordance with the Declaration of Helsinki principles, and was approved by the Georgetown University Institutional Review Board (IRB #2015-1048). All participants provided written informed consent prior to inclusion in this study, and were identified by numerical codes thereafter. Thirty-seven patients with clinically definitive multiple sclerosis were included in this study. Of these patients, 11 were characterized as having active relapsing-remitting MS (aRRMS), nine as having non-active relapsing-remitting disease (naRRMS), and 6 as having secondary progressive MS (SPMS). All patients were non-pregnant, DMD-naïve females aged 18-65 who were being seen by neurologists at Georgetown Multiple Sclerosis and Neuroimmunology Center (GMSNC) and were residents of the Washington D.C. metropolitan area. MS diagnosis and clinical subtype was defined according to the revised McDonald criteria (2010 revisions^16^), and Lublin and Reingold clinical course definitions (2013 revisions^17^). Specifically, aRRMS phenotype was defined as having clinically definitive RRMS with a radiological or clinical relapse within 12 months. naRRMS subtype consisted of individuals with clinically definitive RRMS who had not experienced a relapse or radiological changes within the last 12 months. Patients diagnosed with SPMS had not experienced relapses in the past 12 months and exhibited accumulation of disability, defined by a worsening score on the *Kurtzke Expanded Disability Status Scale* (EDSS)^18^ that persisted for six months or more. Healthy control (HC) subjects were defined as having no history of neurological pathology or autoimmune disease, and satisfied the same inclusion/exclusion criteria as their patient counterparts. Commercially available HC samples (Precision Medicine) were also used in this study.

### Collection and handling of patient samples

Patient blood was drawn and obtained from GMSNC or Georgetown University’s Clinical Research Unit (GU-CRU). Three 8ml CPT-citrate vacutainers (BD cat. 362761) were drawn per patient. PBMCs were isolated from CPT-citrate tubes by density gradient centrifugation at 1500*g*, for 30 min at room temperature (RT). Supernatant was removed and pellets were resuspended in PBS and cell count was obtained to make ∼1×10^6^ cells per 2ml cryovial aliquots (ThermoFisher) and stored at -80° for 24h before transfer to liquid nitrogen.

### PBMC culture and T-cell activation

PBMCs were thawed and plated following relevant sections contained the Helmholtz Zentrum Munich Clinical Cooperation Group Immune Monitoring Protocol. Cells were thawed quickly in 37°C water bath, rinsed twice, centrifuged and resuspended in 10ml 37° C.T.L. Test medium (ImmunoSpot cat. CTLT-005 plus 1% GlutaMAX™ and 1% penicillin-streptomycin). Total live cell count was performed with Trypan Blue (Sigma). Cell density was brought to 1×10^6^ cells/ml in test medium, and plated as 900μl cell suspension per well of a 12-well culture plate. For T-cell stimulation and treatment, plated cells were immediately supplemented with ImmunoCult™ human CD3/CD28/CD2 T cell activator tetrameric antibody complex (Stemcell Technologies cat. 10970) at 20μl/ml, or PBS, and incubated at 37°C for 48h, followed by addition of recombinant human IL4I1 protein (200ng/ml) (R&D cat. # 5684-AO) or 1XPBS (sham-treatment). After 24h incubation, cells were prepared for qRT-PCR analysis or for engrafting into athymic mouse lesions.

### cDNA synthesis and qRT-PCR

Total RNA was extracted from cells with TRIzol™. High integrity RNA (RIN > 7) was used to synthesize cDNA with iScript™ gDNA Clear cDNA Synthesis Kit (Bio-Rad). PCR primers for human *B-ACTIN, IFNG, IL10, IL17A, IL4I1, TGFβ, TNFα*, and *TOB1* were purchased from Bio-Rad. Sybr-Green RT-PCR was performed using the SsoAdvanced Universal SYBR® Green Supermix (Bio-Rad) and analyzed by the CFX96 Touch™ Real-Time PCR Detection System (Bio-Rad). All samples were assayed in triplicate. Results were normalized against *B-ACTIN* and expressed as mean ± SEM.

### Flow cytometry

Flow cytometric analysis was performed on healthy control PBMCs after 48h stimulation with ImmunoCult™ human CD3/CD28/CD2 T cell activator tetrameric antibody complex (Stemcell Technologies cat. 10970) at 20μl/ml, or PBS (sham-stimulation) and 24h treatment with recombinant human IL4I1 protein (R&D cat. 5684-AO) or 1XPBS. All protocols and reagents used were designed and purchased from BioLegend. Protein transport inhibitor Brefeldin A was included in the last 4 hours of cell culture activation in order to block cytokine transport. Then the cells were stained with CD4, CD8, IL10, IFNγ, L17, CD56, CD14 and CD15 primary antibodies following the BioLegend intracellular cytokine staining protocol. Stained cells were analyzed at the Georgetown Lombardi Comprehensive Cancer Center Flow Cytometry & Cell Sorting Shared Resource (FCSR). CD3 antibody was not used because the cells were treated with anti-CD3/CD28/CD2 for activation. Cells that were negative for CD65/CD14/CD15 were considered as CD3+. For the T helper cell population, cells were gated on Live/Dead and CD3+; analyzed for CD4 and CD8. For CD4+ T helper cell subtypes, cells were gated on Live/Dead, CD3+, CD4+; analyzed for IFNγ, L17, and IL10.

### Mouse demyelination and lymphocyte grafting

All animal experiments were performed in accordance with the ARRIVE (Animal Research: Reporting In Vivo Experiments) guidelines, and in accordance with approved Institutional Animal Care and Use Committee (IACUC) protocols of Georgetown University. Female athymic, “nude” mice (Foxn1^nu^) were purchased from Jackson Laboratory (Bar Harbor, ME). Focal demyelination was induced by injecting 1.0% lysolecithin (Sigma) in 1XPBS into the spinal cord dorsal horn of female athymic mice at 8-10 weeks old. PBMCs from healthy control donors were stimulated with anti-CD3/CD28/CD2, and treated with IL4I1 or PBS. Treated cells were collected in media, centrifuged at 600 x g for 10min at 4°C, and resuspended in 1XPBS at 10^5^ live cells/μl. 1μl PBMCs suspension was injected into the lesion site, which was previously marked by charcoal, at 48 hours after lysolethicin injection. Non-grafted control mice (NGC) did not undergo the grafting procedure. Mice in each of 3 groups (NGC, PBS-grafted and IL4I1-grafted), with n=3-5 per group, were sacrificed 15 days after the lysolethicin-injection was performed.

### Tissue sectioning and immunohistochemistry (IHC)

Mice were perfusion-fixed with 4% (w/v) paraformaldehyde (PFA). Spinal cord tissues were dissected and postfixed for 45 min in 4% PFA at RT. Tissues were cryoprotected in 20% (w/v) sucrose before freezing in OCT, and cryosectioned at 12 μm on SuperFrostPlus slides (Stellar Scientific), and stored at −80°C until use. Sections were thawed for 30 min at RT and incubated in blocking solution (0.1% [v/v] Triton X-100 and 10% fetal bovine serum in TBS) for 1 hour at RT. Rat anti-myelin basic protein (MBP) primary antibody (AbD Serotec) was diluted 1:400 in TBS blocking solution and applied to sections overnight at 4°C. AlexaFluor® 488 secondary antibody (ThermoFisher) was used at a concentration of 1:500 and applied for 45 min at RT. FluoroMyelin™ dye (ThermoFisher) at was applied for 45 min at 4°C, at a concentration of 1:75 in TBS blocking solution. CC1 mouse antibody (Millipore), at a concentration of 1:100 in blocking buffer, and Olig2 rabbit (Millipore), at a concentration of 1:500 in blocking buffer, were added on tissues for 48 hours at 4°C after Mouse-On-Mouse Detection (Vector Laboratories). To label nuclei, Hoechst stock (ThermoFisher) was diluted 1:20000 in TBS and applied with secondary antibodies for 45 min.

### Imaging and quantification of IHC

For quantification of IHC, stained areas were manually captured under the LSM880 confocal microscope, and a minimum of 3 sections from n=3-5 mice were examined. ImageJ software was used to quantify MBP intensity. Images were de-identified such that analyses were blinded to experimental conditions. Demyelinated lesions were identified as areas of high cellular density, as defined by nuclear (DAPI) staining. Fluorescence intensity in non-lesioned areas was measured on each channel in order to correct for background fluorescence. Corrected Fluorescence Intensity (CFI) was calculated as: *CFI = IntDensity of lesion – (area of lesion* x *mean fluorescence of background*) per channel. CFIs were averaged per mouse per condition and transferred to GraphPad Prism 7 (La Jolla, CA, USA) for normalization and graphic representation. For Olig2/CC1 quantifications, lesion areas were identified based on nuclei density, and Olig2/CC1 positive cells were quantified and lesion area assessed via ImageJ software.

### Statistics

All statistics were performed using GraphPad Prism 7 (La Jolla, CA, USA). Data is represented as mean ± SEM after identifying statistical outliers. Student’s *t* tests were conducted for paired, within-group analyses; ordinary one-way ANOVA with Tukey’s multivariable comparisons *post-hoc* analysis (alpha=0.05) are used for unpaired, between-group comparisons. Statistical significance is reported as ns, *P ≤ 0.05, **P ≤ 0.01, ***P ≤ 0.001, ****P ≤ 0.0001.

## RESULTS

### IL4I1 skews CD4^+^ T cell populations from healthy donors towards a regulatory state

To examine the effect of IL4I1 on human lymphocytes, peripheral blood mononuclear cells (PBMCs) were isolated from healthy donor blood samples and activated with anti-CD3/CD28/CD2 microbeads in culture. After 48 hours of stimulation, recombinant human IL4I1 or PBS (vehicle control) were added for 24 hours, followed by flow cytometry analysis. We found that that the relative percentages of CD4^+^ and CD8^+^ T-cells were similar between IL4I1 and PBS treatment, suggesting that IL4I1 does not influence CD4^+^ or CD8^+^ T-cell polarization (**Fig. 1A**). Among the CD4^+^ T-cell populations, we found that IL4I1 increased the relative percentage of regulatory T-cells (Tregs cells: CD4^+^ IL10^+^ IFNγ^−^) by over five-fold (3.48% *vs*. 0.66%), and Th2 cells (CD4^+^ IL17^−^ IFNγ^−^ IL10^−^) by nearly two-fold (67.18% *vs*. 37.16%) compared to control (**Fig. 1B**). Furthermore, pro-inflammatory Th17 (CD4^+^ IL17^+^ IFNγ^−^) populations were reduced substantially in IL4I1-treated samples by over thirty-fold (0.96% *vs*. 30.80%) (**Fig. 1B**). The modulatory effect of IL4I1 on human T cells supports our previous findings in mouse models of MS,^11^ therefore suggesting that IL4I1 regulates CD4^+^ T-cell function.

**Figure 1.**
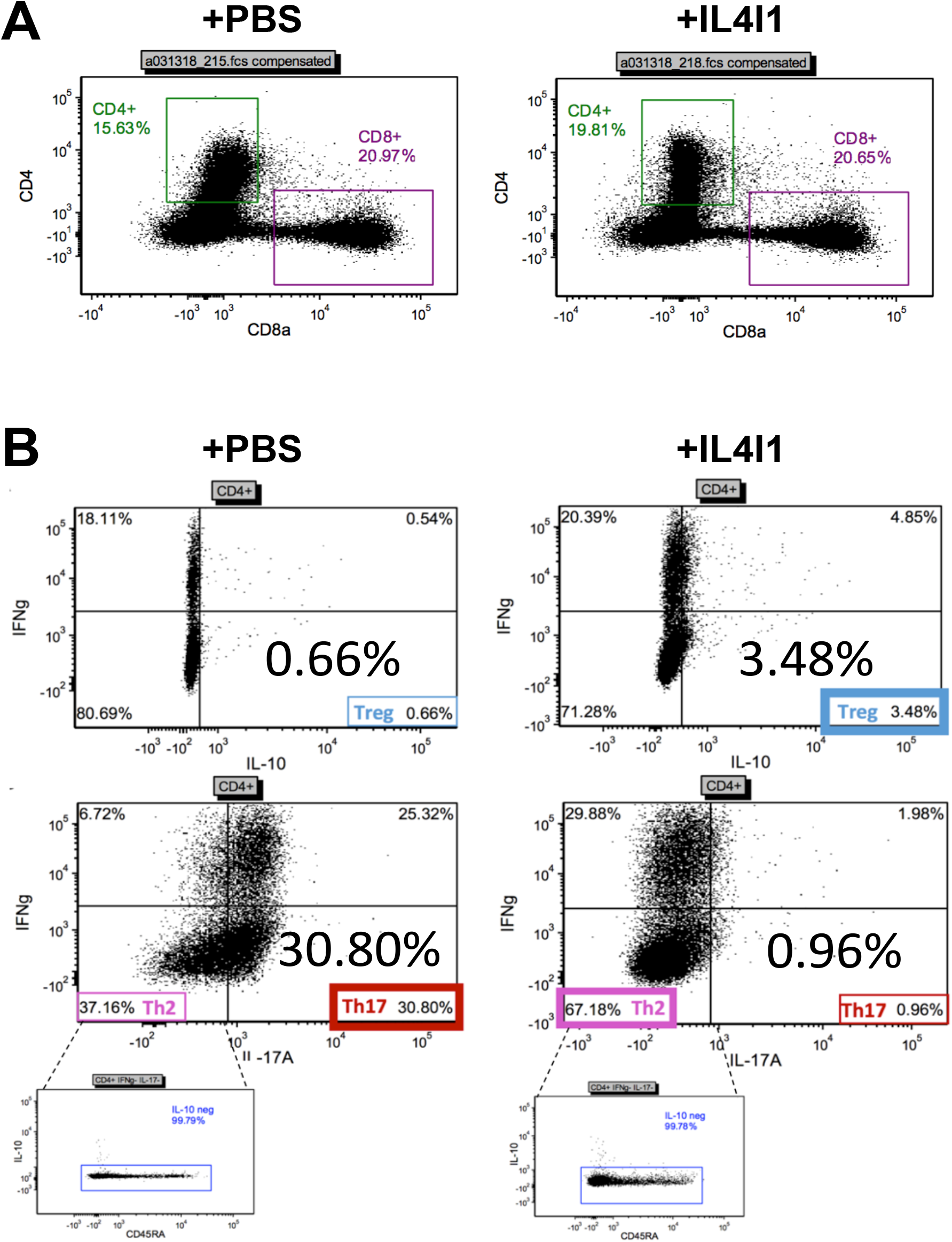
IL4I1 increases Treg and Th2 cell densities, and decreases Th17 cell density in PBMC-derived lymphocytes. Flow cytometry analysis of PBMC-derived lymphocytes from healthy control donor under untreated (+PBS) and treated (+IL4I1) conditions. (**A**) Analysis of CD4^+^ *vs*. CD8^+^ cells. (**B**) Percentage of regulatory T cell (Treg), type 2 helper T cell (Th2), T helper 17 cell (Th17) subtypes among CD4^+^ T-cells based on IFNγ *vs*. IL10 and IFNγ *vs*. IL17 expressions.

### Remyelination is enhanced in mouse lesions engrafted with IL4I1-treated human lymphocytes

A recent study by El Behi and colleagues demonstrated that human lymphocytes from MS patients significantly impairs remyelination efficiency in athymic nude mice (*Foxn1*^*nu*^) when engrafted directly into demyelinated lesions.^19^ To determine if the effect of IL4I1 on healthy human lymphocytes affects CNS remyelination *in vivo*, we grafted IL4I1-or vehicle-treated human lymphocytes (1×10^5^) into focally demyelinated lesions of *Foxn1*^*nu*^ mouse spinal cord at 2 days post lesion (dpl). Additionally, CNS lesions without cell engraftment were examined as control. To quantify the relative extent of remyelination, FluoroMyelin™ staining and myelin basic protein (MBP) immunostaining were performed at 15 dpl, corresponding with oligodendrocyte differentiation and early remyelination,^20^ and the relative fluorescence intensities within lesions were examined. We found that lesions engrafted with IL4I1-treated cells exhibited greater FluoroMyelin™ and MBP staining intensity compared to those engrafted with vehicle-treated cells (**Fig. 2A** and **B**). Moreover, MBP staining in lesions with vehicle treated grafts were not significantly different from lesions without any cells engrafted. Quantification of staining intensity revealed that engraftment of IL4I1-treated cells significantly increased FluoroMyelin™ labeling in lesions by two-fold (**Fig. 2B**). To confirm that increased myelin staining resulted from increased oligodendrocytes in lesions, co-immunostaining analysis for the oligodendrocyte lineage cell marker Olig2 and the mature oligodendrocyte marker CC1 was performed. We found that lesions that received IL4I1-treated grafts exhibited significantly greater number of Olig2^+^CC1^+^ oligodendrocytes compared to PBS-treated grafts, and to those without grafts (**Fig. 2C**). Moreover, no difference in oligodendrocyte density was observed between non-grafted control lesions and lesions containing PBS-treated grafts, supporting previous finding^19^. These results suggest that lymphocyte presence alone does not affect remyelination in lesions, and that IL4I1 modulation of CD4^+^ T-cells is necessary to promotes oligodendrocyte remyelination.

**Figure 2.**
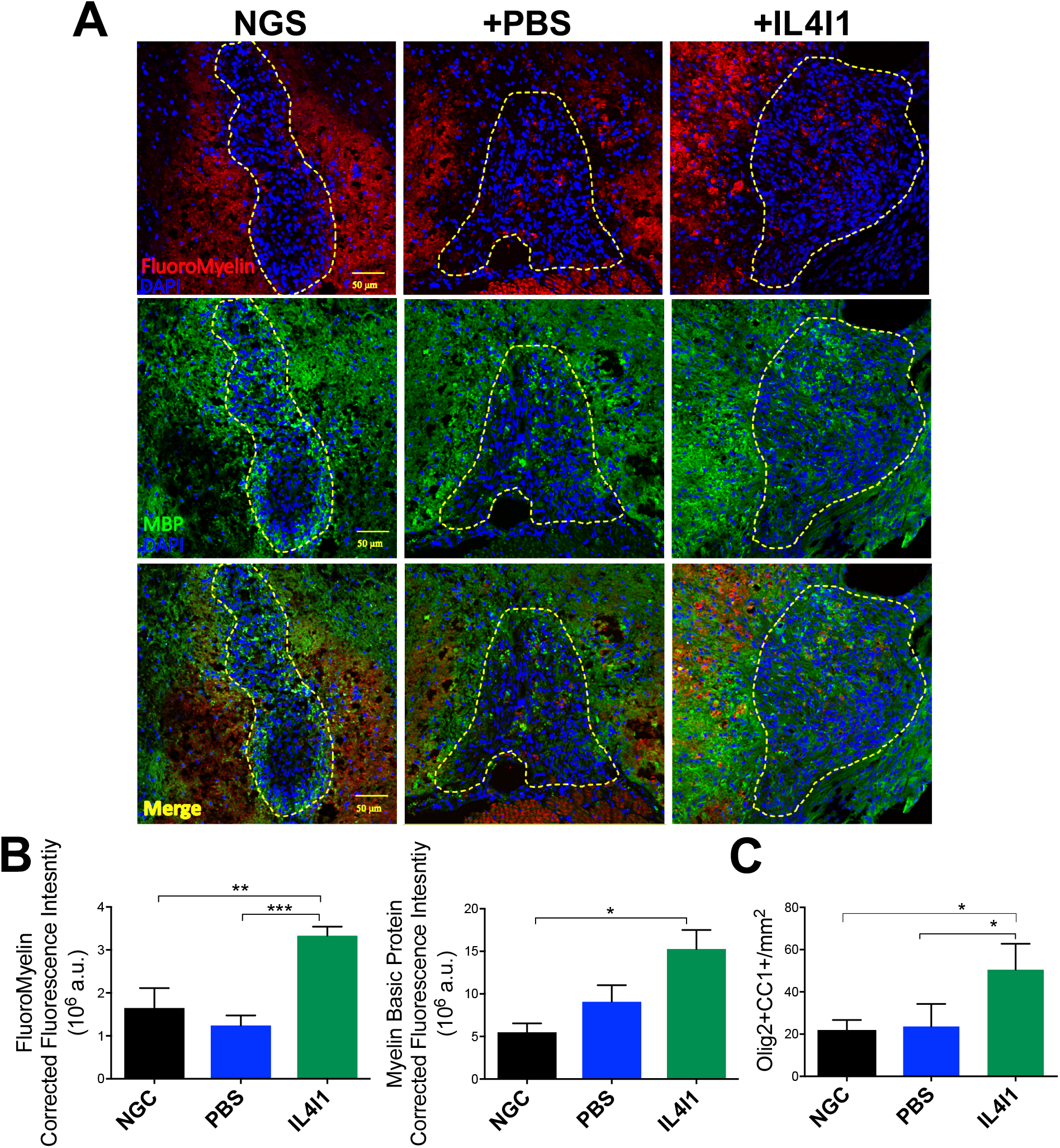
IL4I1-treated human lymphocyte grafts enhance remyelination in nude mouse demyelinated lesions. IHC staining and quantification of myelin within spinal cord dorsal horn lesions of nude mice at 15 days-post lesion (dpl). At 2dpl, lesions were engrafted with IL4I1-treated human lymphocytes (IL4I1; n=4), sham-treated human lymphocytes (PBS; n=5) or were used as non-grafted controls (NGC; n=3). Representative 20x images showing (**A**) FluoroMyelin dye, myelin basic protein (MBP) stain, and merged staining in NGC, PBS and IL4I1 lesions. Lesioned areas are outlined in yellow. Quantification of (**B**) FluoroMyelin and MBP fluorescence intensities, and (**C**) Olig2^+^CC1^+^ cell densities within NGC, PBS-grated and IL4I1-grafted lesions. Scale bar, 50 μm. One-way ANOVA with Tukey’s multiple comparisons test; Data represented as mean ± SEM; *P<0.05, **P<0.01, ***P<0.001.

### Effect of IL4I1 on PBMCs of MS-derived lymphocytes

The observation that IL4I1 modulates T-cells and promotes remyelination raises the question whether endogenous IL4I1 expression or function is impaired in MS. To answer this question, we examined the expression of *IL4I1* mRNA expression in non-activated PBMCs of healthy control (HC) and MS patients. Baseline qRT-PCR analysis of HC-derived PBMCs confirmed that *IL4I1* is expressed at detectable levels (**Fig. 3A**) To identify potential correlation between MS disease state and *IL4I1* levels, *IL4I1* mRNA expression in PBMCs from active relapsing remitting MS (aRRMS), non-active relapsing remitting MS (naRRMS), and secondary progressive MS (SPMS) were compared with HC (see Methods for MS classifications). We found that *IL4I1* expression was significantly reduced in naRRMS compared to HC, with aRRMS and SPMS trending towards reduced levels (**Fig. 3B**). These findings suggest that *IL4I1* is reduced in some MS disease stages.

**Figure 3.**
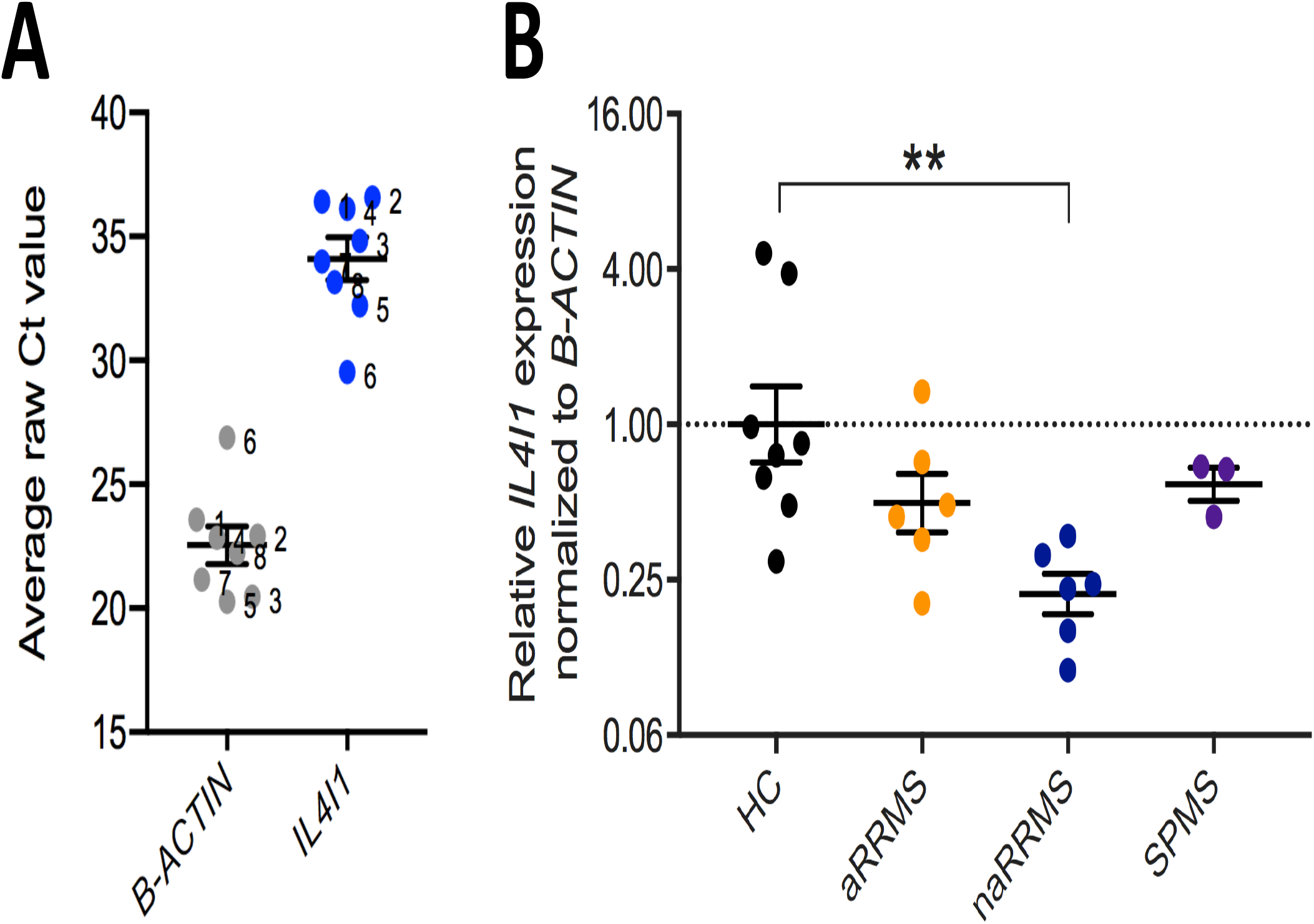
Baseline *IL4I1* expression is reduced in MS patient PBMCs. (**A**) *IL4I1* mRNA expression in unstimulated PBMCs from HC by qRT-PCR. *B-ACTIN* served as a loading control. qRT-PCR Ct values are shown with numbered dots representing individual donors (n=8). (**B**) Relative *IL4I1* mRNA expression in unstimulated PBMCs from HC (n=8) vs. patients with MS. Female donors with aRRMS (n=6), naRRMS (n=6), and SPMS (n=3) are shown relative to control. One-way ANOVA with Tukey’s multiple comparisons test; Data represented as mean ± SEM; **P<0.01.

To examine the effect of IL4I1 on inflammatory cell activity in the context of MS and non-disease states, lymphocytes from HC, aRRMS, naRRMS and SPMS groups were treated with IL4I1 or PBS (control) followed by qRT-PCR analysis for *IL17* expression, since Th17 cells from healthy lymphocytes were most profoundly affected by IL4I1 treatment (**Fig. 1B**). We found that IL4I1 treated HC lymphocytes exhibited reduced *IL17* expression relative to PBS control (**Fig. 4A**). However, its expression was unaffected by IL4I1 in all of the MS derived lymphocytes tested (**Fig. 4A**). This result suggests that IL4I1 is unable to modulate Th17 cells in MS despite its ability to alter Th17 cell activity in healthy patients. Since previous study suggested that IL4I1 targets *TOB1*, an anti-proliferative regulator to prevent Th17 cell expansion,^14^ we next examined the expression of *TOB1* in HC- and MS-derived lymphocytes after IL4I1 or PBS (control) treatment. We found that IL4I1 increased *TOB1* expression in HC lymphocytes compared to control (**Fig. 4B**). However, IL4I1 had no effect on *TOB1* expression in MS lymphocytes in all the disease stages examined (**Fig. 4B**). Collectively, these data suggest that IL4I1 affects *IL17* and *TOB1* expression in healthy lymphocytes, but this relationship is perturbed in MS. Moreover, these results suggest that MS patients exhibit impaired IL4I1-TOB1 signaling that may lead to uncontrolled Th17 cell proliferation.

**Figure 4.**
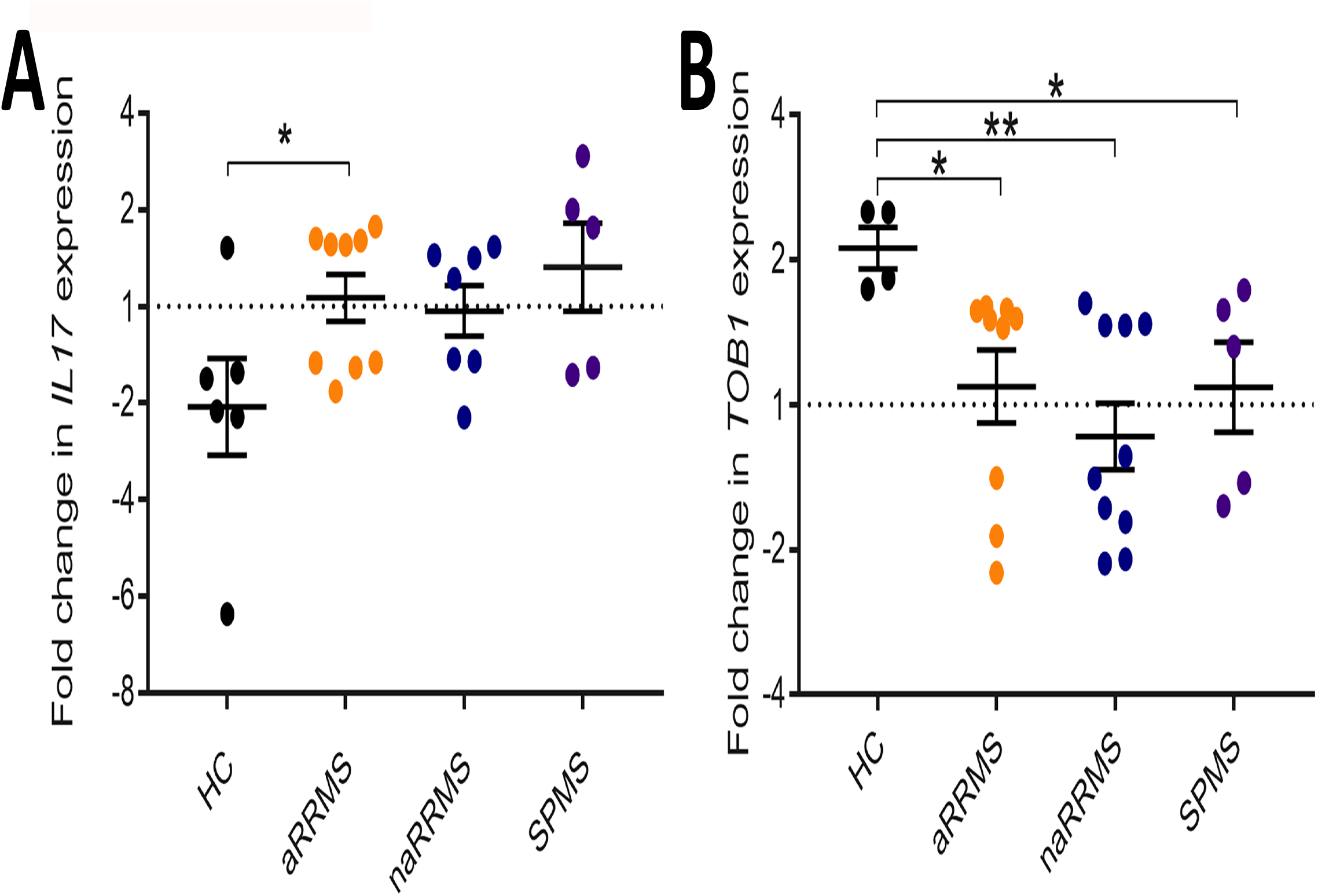
IL4I1 does not affect *IL17* or *TOB1* expression in MS lymphocytes. Fold change of (**A**) *IL17* and (**B**) *TOB1* in PBMC-derived lymphocytes from HC (n=6), aRRMS (n=9), naRRMS (n=6), and SPMS (n=5) after treatment with IL4I. Relative expressions were normalized to their respective PBS-treated controls. One-way ANOVA with Tukey’s multiple comparisons test; Data represented as mean ± SEM; *P<0.05, **P<0.01.

## DISCUSSION

In this study, we have found that IL4I1 skews lymphocytes from healthy individuals towards a regulatory state by reducing the population of Th17 cells and increasing the densities of Tregs and Th2 cells. This shift in lymphocyte subtypes appears to be crucial for promoting for CNS repair. Recent studies have shown that lymphocyte-derived mediators facilitate oligodendrocyte differentiation through interactions with microglial cells, and that Tregs play a critical role in remyelination success in mice.^19,21^ Remarkably, we found that the *in vitro* effect of IL4I1 on healthy human lymphocytes resulted in significantly enhanced remyelination *in vivo* when IL4I1-treated lymphocytes were transplanted into lesions in nude mouse. This result suggests that IL4I1 modulates the inflammatory microenvironment driven by T-cells, thereby promoting remyelination. Additionally, by grafting human lymphocytes into nude mouse CNS lesions, we were able to differentiate between the contributions of graft *vs*. host immune cells and observe the direct impact of human lymphocyte activity on CNS remyelination.^19^ Since PBMCs can be easily obtained from patients, this strategy may serve as a predictive preclinical assay to determine whether immunomodulatory drugs can improve remyelination in MS.

We also found that lymphocytes derived from MS patients exhibited lower expression of *IL4I1* compared to lymphocytes from healthy controls, indicating the transcriptional regulation of *IL4I1* might be perturbed in MS. Since the *IL4I1* genomic locus is situated in a chromosomal region implicated in autoimmune susceptibility,^13^ our results support the potential dysregulation of *IL4I1* in MS. While treatment of lymphocytes from healthy controls with IL4I1 resulted in altered *IL17* and *TOB1* expression, we observed no effect on their expression in lymphocytes from MS patients. These findings suggest that that IL4I1 modulates CD4^+^ T-cell function in healthy human lymphocytes, but is unable to exert its influence on lymphocytes in MS. It is possible that T-cells in MS are resilient or no longer responsive to IL4I1, which may contribute to aberrant inflammation and remyelination impairment in MS lesions. Therefore, future studies aimed at understanding how IL4I1 dysregulation impacts T-cell function in MS would greatly improve our understanding of MS pathogenesis, and potentially lead the development of treatments targeting IL4I1 signaling to promote repair in MS.

## AUTHOR CONTRIBUTIONS

S.E.D., F.S.A., A.W. and J.K.H. initiated and designed this study. S.E.D. conducted *in vitro* and *in vivo* experiments, performed gene expression analysis, analyzed data, and drafted the manuscript for intellectual content. J.H. prepared samples for flow cytometry analysis. S.E.N. and M.N.K. assisted with spinal cord sectioning and performed immunostaining. M.B. and S.E.N. assisted in perfusions and contributed to data analyses.

H.O. assisted with *in vitro* experiments. F.S.A. oversaw patient recruitment and assisted in blood collection. A.W. contributed to qPCR and PBMC analysis. J.K.H. drafted the manuscript for intellectual content, and oversaw the project.

## STUDY FUNDING

This project was supported in part by funding from the NIH (5R01NS107523) and National Multiple Sclerosis Society (NMSS) Harry Weaver Neuroscience Scholar Award (JF-1806-31381) to J.K.H., TurnFirst Foundation to S.E.D. and J.K.H., and NIH-NCATS Training Award TL1TR001431 to S.E.D.

## ACKNOWLEDGEMENT

We thank the Georgetown University Flow Cytometry & Cell Sorting Shared Resource (FCSR), and Tissue Culture Shared Resource (TCSR) for assistance in flow cytometry and PBMC collection, BioLegend for guidance on flow cytometry, Dr. Konstantina Psachoulia for guidance on *in vivo* experimental setup and analysis, Drs. Eveline Vietch and Bevan Main for guidance on qRT-PCR analysis, and members of the Huang lab for helpful discussion and suggestions on this project.

## DATA AVAILABILITY

The datasets used and/or analyzed during the current study available from the corresponding author on reasonable request.

## Notes

**Conflict of interest statement:** The authors have declared that no conflict of interest exists.

### Competing Interest Statement

The authors have declared no competing interest.

## REFERENCES

1. Dutta R, Trapp BD. Relapsing and progressive forms of multiple sclerosis: insights from pathology. Curr Opin Neurol. 2014;27:271–278.

2. Compston A, Coles A. Multiple sclerosis. Lancet. 2002;359:1221–1231.

3. Reich DS, Lucchinetti CF, Calabresi PA. Multiple Sclerosis. N Engl J Med. 2018;378:169–180.

4. Dutta R, Trapp BD. Mechanisms of neuronal dysfunction and degeneration in multiple sclerosis. Prog Neurobiol. 2011;93:1–12.

5. Franklin RJM, ffrench-Constant C, Edgar JM, Smith KJ. Neuroprotection and repair in multiple sclerosis. Nat Rev Neurol. 2012;8:624–634.

6. Fitzner D, Simons M. Chronic progressive multiple sclerosis - pathogenesis of neurodegeneration and therapeutic strategies. Curr Neuropharmacol. 2010;8:305–315.

7. Franklin RJM. Why does remyelination fail in multiple sclerosis? Nat Rev Neurosci. 2002;3:705–714.

8. Compston A, Coles A. Multiple sclerosis. Lancet Lond Engl. 2008;372:1502–1517.

9. Yue Y, Huang W, Liang J, et al. IL4I1 Is a Novel Regulator of M2 Macrophage Polarization That Can Inhibit T Cell Activation via L-Tryptophan and Arginine Depletion and IL-10 Production. PloS One. 2015;10:e0142979.

10. Carbonnelle-Puscian A, Copie-Bergman C, Baia M, et al. The novel immunosuppressive enzyme IL4I1 is expressed by neoplastic cells of several B-cell lymphomas and by tumor-associated macrophages. Leukemia. 2009;23:952–960.

11. Psachoulia K, Chamberlain KA, Heo D, et al. IL4I1 augments CNS remyelination and axonal protection by modulating T cell driven inflammation. Brain. 2016;139:3121–3136.

12. Becker KG, Simon RM, Bailey-Wilson JE, et al. Clustering of non-major histocompatibility complex susceptibility candidate loci in human autoimmune diseases. Proc Natl Acad Sci U S A. 1998;95:9979–9984.

13. Chavan SS, Tian W, Hsueh K, Jawaheer D, Gregersen PK, Chu CC. Characterization of the human homolog of the IL-4 induced gene-1 (Fig1). Biochim Biophys Acta. 2002;1576:70–80.

14. Santarlasci V, Maggi L, Mazzoni A, et al. IL-4-induced gene 1 maintains high Tob1 expression that contributes to TCR unresponsiveness in human T helper 17 cells. Eur J Immunol. 2014;44:654–661.

15. Corvol J-C, Pelletier D, Henry RG, et al. Abrogation of T cell quiescence characterizes patients at high risk for multiple sclerosis after the initial neurological event. Proc Natl Acad Sci U S A. 2008;105:11839–11844.

16. El Behi M, Sanson C, Bachelin C, et al. Adaptive human immunity drives remyelination in a mouse model of demyelination. Brain. 2017;140:967–980.

17. Jeffery ND, Blakemore WF. Remyelination of mouse spinal cord axons demyelinated by local injection of lysolecithin. J Neurocytol. 1995;24:775–781.

18. Dombrowski Y, O’Hagan T, Dittmer M, et al. Regulatory T cells promote myelin regeneration in the central nervous system. Nat Neurosci. 2017;20:674–680.

19. Polman CH, Reingold SC, Banwell B, et al. Diagnostic criteria for multiple sclerosis: 2010 revisions to the McDonald criteria. Ann Neurol. 2011;69:292–302.

20. Lublin FD, Reingold SC, Cohen JA, et al. Defining the clinical course of multiple sclerosis: the 2013 revisions. Neurology. 2014;83:278–286.

21. Kurtzke JF. Rating neurologic impairment in multiple sclerosis: an expanded disability status scale (EDSS). Neurology. 1983;33:1444–1452.

